# GM1 asymmetry in the membrane stabilizes pores

**DOI:** 10.1101/2022.01.21.477228

**Authors:** Mina Aleksanyan, Rafael B. Lira, Jan Steinkühler, Rumiana Dimova

## Abstract

Cell membranes are highly asymmetric and their stability against poration is crucial for survival. We investigated the influence of membrane asymmetry on electroporation of giant unilamellar vesicles with membranes doped with GM1, a ganglioside asymmetrically enriched in the outer leaflet of neuronal cell membranes. Compared to symmetric membranes, the lifetimes of micronsized pores are about an order of magnitude longer suggesting that pores are stabilized by GM1. Internal membrane nanotubes caused by the GM1 asymmetry, obstruct and additionally slow down pore closure, effectively reducing pore edge tension and leading to leaky membranes. Our results point to the drastic effects this ganglioside can have on pore resealing in biotechnology applications based on poration as well as on membrane repair processes.

**SIGNIFICANCE:** Membrane pore closure is crucial for cell survival and is important for biotechnological and medicine applications based on transfer of material, e.g. drugs, genes, through pores. Electroporation is widely used as means to perforate the membrane but factors governing membrane resealing are still a matter of debate, in particular the large variations of pore lifetimes. Here, we probed the effect of bilayer asymmetry on pore dynamics employing cell-sized giant unilamellar vesicles doped with the ganglioside GM1 (asymmetrically enriched in neurons). We find that the presence of GM1 and its asymmetric distribution in the bilayer dramatically slows down pore resealing, not only by mere molecular stabilization of the pore rim, but also by generating membrane nanotubes.

## INTRODUCTION

Pores in membranes allow for exchange and introduction of substances in cells, and when generated exogenously, their closure is crucial for cell survival. Among the different approaches of generating pores (1), electroporation offers the attractive feature of precise localization and temporal control. Thus, electroporation-based techniques have gained significant importance in biotechnology over the years as one of the low-cost, safe, practical and efficient ways of accessing the inner compartments of cells and controlling biological processes (2-5). Depending on the duration, strength and number of electric pulses, membrane electroporation can either result in a permanent cell lysis (i.e. irreversible electroporation), or temporary poration followed by membrane resealing (i.e. reversible electroporation) (4). Irreversible electroporation has broad applications in regenerative medicine (6) and tissue engineering (7) including tumor ablation (8), electrochemotherapy (9,10) and exogenous cell engraftment (11). On the other hand, reversible electroporation has been one of the most widely applied methods in biomedical engineering (12) including anticancer treatment (13), gene and drug delivery (14), cell transfection (15) and inactivation of microorganisms (16). Electroporation thresholds and pore kinetics are known to differ from cell to cell and depend on a large variety of parameters including pulse shape, duration, number and repetition, cell size and state as well as environmental conditions (17). Despite the numerous theoretical and experimental studies on electroporation (see e.g. refs. (18-20)), the fundamental mechanisms underlying the plasma membrane resealing after an electrical breakdown have not been yet fully explained; for example, it remains unclear why pores have a wide range of lifetimes spanning from milliseconds to minutes (21-23). However, understanding the detailed membrane reorganization and pore stability is crucial for the optimization of clinical settings of electroporation as well as membrane repair and wound healing.

To resolve the underlying mechanisms of poration and stability of plasma membranes, giant unilamellar vesicles (GUVs) (24,25) as cell-size simple membrane systems are commonly investigated (26-28) because the membrane response is directly accessible via optical microscopy. Electroporation is initiated by the increase in membrane tension induced by the electric field. Above the membrane electroporation threshold, pores are formed, relaxing the tension (29,30). The pores can spontaneously reseal driven by membrane edge tension (31-33), the energetic cost of lipid rearrangement along the pore rim. Not surprisingly, the edge tension depends on membrane composition as well as on the medium (presence of ions, molecules or detergents) (34-36).

The response of single-component (symmetric) GUVs to electric fields has been thoroughly explored (26,27,30). For such simple membranes, the application of a single DC pulse can lead to GUV deformation and formation of micron-sized pores (macropores). The lifetime of these pores is on the order of a few hundreds of milliseconds (30,37). However, very simple model membranes might not sufficiently well represent the response of the complex plasma membrane, which exhibits both sophisticated composition and leaflet asymmetry. Here, we explore the effect of asymmetry albeit in a simple model membrane, namely one made of palmitoyloleoylphosphocholine (POPC) and doped with the ganglioside GM1. GM1 is involved in many biological events as one of the major components of the outer leaflet of the mammalian membranes (38,39). In addition, it is asymmetrically distributed and abundant in the nervous system and is associated with neuronal differentiation and development processes (40). GM1 asymmetry in GUV membranes was found to induce substantial membrane curvature, leading to the formation of membrane nanotubes (41,42). In addition, the lifetime of electro-induced pores in GM1-doped GUVs was found to be orders of magnitude longer than pores formed on typical (and symmetric) POPC GUVs (30). This naturally raises the question of whether it is just the presence of GM1 or also its asymmetric distribution in the membrane that dramatically slows down pore closure. Answering this question can shed light on the stability of cells when porated (not only via electric fields) and help resolve mechanisms of plasma membrane repair. Thus, we aimed at investigating in detail the resealing dynamics of electroporated GM1-doped GUVs both as a function of GM1 fraction and membrane asymmetry.

## MATERIALS AND METHODS

The phospholipid 1-palmitoyl-2-oleoyl-sn-glycero-3-phosphocholine (POPC), the fluorescent lipid analogue 1,2-dipalmitoyl-sn-glycero-3-phosphoethanolamine-N-(lissamine rhodamine B sulfonyl) (DPPE-Rh) and the ganglioside GM1(Ovine Brain) were purchased from Avanti Polar Lipids (Alabaster, AL). Glucose, 0.5 Na HEPES and the fluorescent dye calcein were purchased from Sigma Aldrich (St. Louis, MO USA). Stock solutions of the phospholipid, the dye and the ganglioside GM1 were prepared in a mixture of dichloromethane:methanol (2:1 volume) and the solutions were stored at -20°C until the usage. All the microscopic observations were done with home-made chambers assembled from 26×56 mm and 22×22 mm cover slips purchased from Thermo Fisher Scientific (Waltham, MA USA). Cover slips were rinsed with ethanol and distilled water before usage. To measure the solution osmolarity, Osmomat 3000 osmometer (Gonotec GmbH, Berlin, Germany) was used.

GUVs were prepared by electroformation method (43), see section S1 in the supporting information (SI) for details. For the generation of GM1 leaflet asymmetry, GM1-doped GUVs were 10-fold diluted in isotonic 1 mM HEPES buffer, see SI section S2. For the electroporation experiments, GUVs were subjected to a single direct current (DC) pulse (0.3 – 0.6 kV/cm, 3 or 50 ms) and their responses were recorded either with confocal microscopy, or under epifluorescence or phase contrast combined with high speed imaging, see SI section S3. The membrane edge tension was deduced from analysis of the pore closure (44), see SI section S6. For the analysis of long-term permeation, GUVs were exposed to a single DC pulse and calcein entry was monitored for 5 minutes. Quantification of GUV leakage was performed through fluorescence intensity analysis, see SI section S7. In all experiments, microscopy images were analyzed either with LASX software or ImageJ. All the datasets were analyzed and plotted with Origin Pro software.

## RESULTS AND DISCUSSION

### Dilution of GM1-doped vesicles results in membrane asymmetry

POPC GUVs doped with 0, 2 or 4 mol% GM1 were prepared in 1 mM HEPES buffer and successful incorporation of GM1 into the membrane was confirmed by binding of CTB-Alexa; see SI section S1 and Fig. S1. The explored GM1 concentrations fall in the range found in neurons (45). As previously reported, GM1 in the bilayer is in dynamic equilibrium with GM1 in the surrounding solution (41). Thus, the GM1 concentration in the membrane can be controlled by the concentration of free GM1 in the bulk. Because flip-flop of GM1 molecules is negligible on the experimental time scales, the GM1 concentration outside the vesicle can be used to control GM1 leaflet asymmetry. In our experiments, the GUVs were 10-fold diluted in GM1-free isoosmolar buffer, which resulted in the desorption of a large fraction of the GM1 lipids from the outer membrane leaflet, increasing the asymmetry compared to the inner leaflet (Fig. 1A). For example, for vesicles prepared with 2 mol% GM1, the dilution step results in ganglioside concentration of 0.47 mol% in the outer leaflet and 1.98 mol% in the inner leaflet as characterized previously (41). A direct consequence of this asymmetry is the generation of spontaneous (preferred) membrane curvature (46,47). In vesicles with excess area (small volume-to-area ratio), this spontaneous curvature stabilizes vesicle morphologies with highly curved membrane nanotubes, as demonstrated previously (41,48,49). In the case of the 10-fold diluted vesicles initially prepared with 2 or 4 mol% GM1, the asymmetric distribution of the ganglioside results in negative spontaneous curvature between around −1/(500 nm) and −1/(200 nm) (41) which stabilizes inward nanotubes, see SI section S2 and Fig. S2. In contrast, symmetric vesicles (no dilution) do not form tubes (Fig. S2). In order to assess the membrane stability and the edge tension of symmetric and asymmetric membranes, we applied electric pulses to GUVs of asymmetric or symmetric membranes at varying GM1 concentrations.

**Fig. 1.**
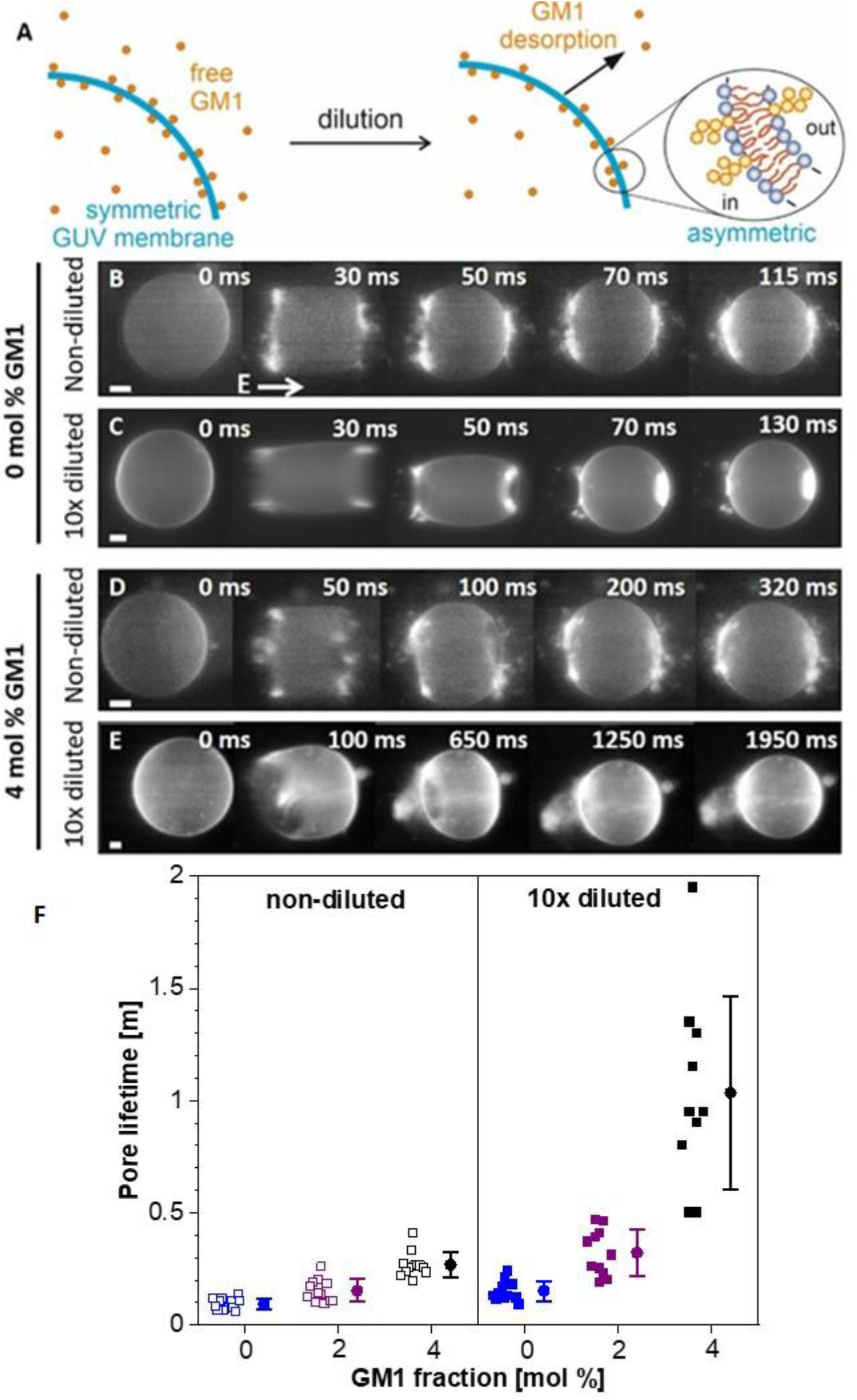
Poration of GUVs with symmetric and asymmetric membranes. (A) Upon dilution, GM1 from the outer GUV leaflet desorbs and renders the membrane asymmetric as illustrated in the sketch. (B-E) Comparison of electroporation of POPC GUVs containing no GM1 (B, C) and 4 mol% GM1 (D, E) in non-diluted external solution (B, D) and upon 10-fold dilution in isotonic external solution (C, E) imaged with epifluorescence microscopy. The membrane was stained with 0.1 mol% DPPE-Rh. The vesicles were exposed to a single DC pulse with amplitude of 0.3-0.4 kV/cm and duration of 50 ms. The direction of the electric field is illustrated with the arrow in (B). The timestamps in the top-right corner of each snapshot show the time after the beginning of the pulse. The sequence in (E) corresponds to Movie S1 in the SI. Scale bars: 10 µm. (F) Macropore lifetimes measured on vesicles (15 to 55 µm in radius) with varying molar fractions of GM1 in non-diluted (open squares) and 10-fold diluted (solid squares) solutions. Every data point indicates a measurement on an individual GUV from all together 3 preparation batches; between 10 and 14 vesicles per composition and condition were explored. Mean values and standard deviations are shown by the solid circles and bars, respectively. Blue, purple and black corresponds to 0, 2, 4 mol% GM1.

### Both GM1 fraction and asymmetry affect pore stability

Pure POPC and 4 mol% GM1-doped GUVs in both non-diluted and diluted solution were exposed to a single, strong electric pulse (50 ms duration, amplitude of 0.3-0.4 kV/cm), Fig. 1; see also SI section S3. Such pulses raise the transmembrane potential above the poration threshold and optically detectable pores (macropores) are created (30). The pore dynamics was monitored via high-speed imaging (SI section S3). POPC GUVs exhibit symmetric membranes in both diluted and non-diluted solutions and the pores developed in these membranes had short lifetimes around 152 ± 45 ms (Fig. 1B,C,F) consistent with previous reports (30,37). On the contrary, non-diluted GUVs symmetrically doped with 4 mol% GM1 exhibited pores with twice longer lifetime on the order of 268 ± 56 ms, Fig. 1D,F.

The above results (Fig. 1) demonstrate that pores are stabilized by GM1 in the membrane. We then investigated the effect of leaflet asymmetry by comparing diluted (asymmetric) and non-diluted (symmetric) GM1-doped vesicles (see sketch in Fig. 1A). Remarkably, in the asymmetric GUVs, as shown in Fig. 1E, pore lifetimes increase dramatically to 1035 ± 432 ms, i.e. 5 times longer than that of symmetric GM1-doped GUVs and 10 times longer than that for pure-POPC membranes (see also Fig. 1F). These findings show that not only the presence of GM1 but also leaflet asymmetry stabilize very long-living membrane pores. It is important to note that for the asymmetric membranes, poration was associated with expelling inward tubes through the macropores, Fig. 1E and SI Movie S1, Fig. S4 and Movie S3. We now discuss three factors for pore stabilization in asymmetric membranes.

(i) Prior to the pulse, inward membrane nanotubes are present in a large fraction of the vesicles (Fig. S2A, B); note that the preparation protocol results in vesicles of different volume-to-area ratio and thus the excess area for tube formation in each vesicle is different. During poration, water flow caused by the higher internal (Laplace) pressure drags nanotubes out through the formed pores (Fig. 2A, SI Movie S2). The tubes protrude from the vesicle interior (Fig. 1E, SI Movie S1) and occasionally also around the pore rim (Fig. 2A, SI Movie S2). Thus, the steric hindrance of the nanotubes on the closing membrane plausibly contributes to increased pore lifetime, also leading to incomplete membrane resealing as shown below.

**Fig. 2.**
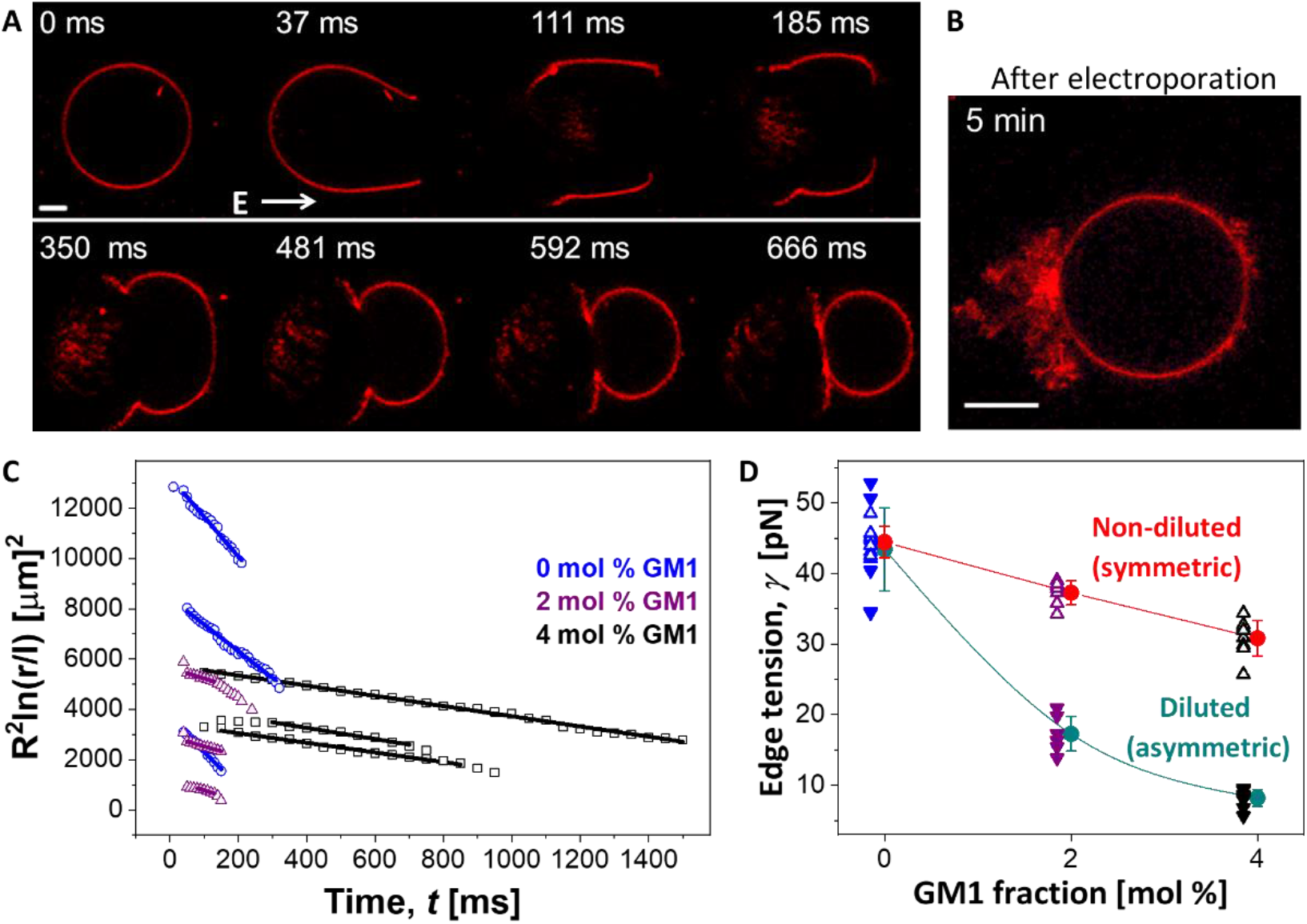
Electroporation and edge tension of symmetric and asymmetric membranes. (A) A typical example of electroporation of a 4 mol% GM1-doped asymmetric GUV observed with confocal microscopy upon application of a DC pulse (0.3kV/cm, 50 ms). The time is relative to the beginning of the pulse. The membrane contains 0.1 mol% DPPE-Rh and the GUV was 10-fold diluted with 1 mM isotonic HEPES buffer. Image sequence corresponds to Movie S2. (B) Cross section of the vesicle 5 minutes after the pulse application. Scale bars: 20 µm. (C) Example datasets showing pore closure dynamics for each of the compositions (three examples per composition) displayed as the porated region *R*^2^*ln*(*r*) versus the time after application of the pulse; where *r* and *R* are the respective pore and vesicle radii. Differences in the absolute values of the data come from the differences in GUV radius *R*. In order to avoid plotting a dimensional value in the logarithmic term, the pore radius is scaled by *l* =1 µm. Open symbols reflect the experimental data (each of which required the manual assessment of the pore radius from 20-40 images) while the solid lines are linear fits, the slope of which yields the edge tension (given in panel D). As the GM1 fraction in the membrane increases, the slope decreases (the slope defines the edge tension *γ*, see SI section S6). At the same time, the pores display longer lifetime. (D) Edge tension versus GM1 fraction for symmetric (open triangles) and asymmetric (solid triangles) GUVs. 6 to 10 vesicles per composition were measured for pure POPC (blue), 2 mol% GM1 (purple) and 4 mol% GM1 (black); same color scheme as in panel C. Mean values and standard deviation of the edge tension are indicated with solid circles and bars for symmetric (red) and asymmetric (green) membranes (see also Fig. S7 for a more detailed display). The curves are a guide to the eyes.

(ii) Pore lifetime is strongly modulated by the spontaneous curvature of asymmetric membranes. A key observation is that during pore opening, in contrast to symmetric POPC GUVs, asymmetric GM1 vesicles exhibit sprouting of new membrane nanotubes, not present prior to the pulse, (SI Figs. S4, S5, Movies S3, S4). This occurs almost immediately after poration, indicating that the outer spherical membrane segment of the vesicle prefers to rearrange into highly bent nanotubes. This process can be understood considering the membrane spontaneous curvature. Symmetric membranes have zero spontaneous curvature, *m* = 0. For the asymmetric membranes *m* is around -1/460 nm^-1^ for vesicles prepared with 2 mol% GM1 and -1/220 nm^-1^ for 4 mol% GM1 (41). Asymmetric vesicles minimize their bending energy by forming cylindrical nanotubes with radius of 1/(2*m*) (50). Before poration (in intact vesicles), the ratio of membrane area stored in tubes to area of the weakly curved outer GUV membrane is set by the osmotically stabilized vesicle volume and total membrane area. However, when the volume constraint is relaxed by membrane poration, resealing of the vesicle pore competes for membrane area with formation of new nanotubes. The latter process reduces the vesicle surface area acting analogously to surface tension (46) that pulls the pore open. Considerations of pore and membrane elastic energy suggest that the asymmetry should result in transforming all available area into nanotubes when a pore of radius *r* > *r*_*c*_ ≡ *γ*/2*κm*^2^ is formed (SI section S5). Introducing the experimentally measured values for the edge tension *γ*, the bending rigidity *κ* and *m* we obtain that pores larger than 15 μm (2 mol% GM1) or 1.4 μm (4 mol% GM1) should become unstable and expand while transforming the membrane into tubes. Typical pore sizes, particularly for the 4 mol% GM1 vesicles were larger and indeed membrane tubulation upon poration was frequently observed for asymmetric GUVs. Almost complete transformation of the vesicle membrane area to a tubular network was also observed occasionally (Fig. S6, Movie S5). However, for most of the vesicles, the period of delayed pore closure and nanotube formation lasts only a few hundred microseconds. Eventually, the membrane pore starts to reseal, which indicates that during pore opening some of the membrane asymmetry is lost, and this corresponds to reduction of the spontaneous curvature *m*. Because of the quadratic relation between critical pore radius *r*_*c*_ and spontaneous curvature *m*, a rather small exchange or loss of GM1 is sufficient to enhance pore closure. One possible mechanism for loss of GM1 asymmetry is the interleaflet exchange of membrane-bound GM1 across the pore edge. Another mechanism leading to suppression of asymmetry (i.e. decrease of *m*, ultimately leading to pore closure) is the desorption of GM1 from the inner vesicle leaflet as the enclosed GUV solution now becomes diluted when the pore opens.

(iii) Additionally, the finite membrane viscosity or steric constraints might limit excessive nanotube formation and complete vesicle destabilization.

In summary, we correlate the long pore lifetimes to steric hindrance by nanotubes protruding through the pore and altered membrane mechanics due to dynamic changes in membrane asymmetry during pore closure.

### Pore edge tension is lowered by GM1

Next, we set to quantitatively explore the dynamics of pore closure in asymmetric and symmetric vesicles and deduce their pore edge tension using a previously reported method (44,51) (for details see SI section S6, Fig. 2C and Fig. S7). The measured edge tension of pure POPC GUVs (Fig. 2D) fall in the range of literature values (35,36) corroborating the consistency of our data and analysis. With increasing GM1 fraction, the mean values of edge tensions obtained from different preparations decrease, see Fig. 2D and SI Table 1. For the asymmetric GM1-doped GUVs, the measured edge tension is an apparent one because of the presence of tubes in the pore area which is not accounted for by the theoretical model (31). The results for symmetric GUVs scale linearly with GM1 fraction; more GM1 causes stabilization of pores and lower values of edge energy.

This effect in symmetric membranes might be understood considering the cone-like shape of GM1, which could favorably locate at the pore rim where the monolayer curvature is high. This molecular effect should be distinguished from the apparent reduction of edge tension by membrane asymmetry detailed above, even though both effects contribute to increasing pore lifetime.

Finally, we return to the membrane remodeling events that were observed during poration of asymmetric GM1 vesicles with nanotubes (Fig. 2A-B, Movies S1, S2). During pore expansion, the electric field and the flow of the solution leaking out orient and extend the tubes out towards the vesicle exterior. As the pore then proceeds to close, it bundles together the protruding tubes resulting in high fluorescence from accumulated membrane material in the form of tubes and small buds at the location of the closed pore. We investigated the influence of this accumulation on the long-term permeability of the vesicles. Here, we distinguish the optically resolvable “macropores” from small “submicroscopic” pores not detected optically.

### Asymmetric vesicles become leaky after macropore closure

To test long-term membrane permeation of the asymmetric membranes, the vesicles were grown in sucrose solution and diluted in isotonic glucose solution, see SI section S7. As a result of the different refractive index of the sugar solutions, the vesicles appear dark on a brighter background when observed under phase contrast microscopy (SI Fig. S8A). During prolonged observations, we noticed that GM1-doped GUVs lost their optical contrast after macropore closure (Fig. S8), indicating exchange of solution between GUV interior and exterior even after macropore closure. We thus explored whether and to what extent GM1 present in the membrane makes the vesicles leaky. To quantitatively monitor membrane permeability, calcein, a small water-soluble and membrane-impermeable dye, was introduced in the external media (at 5 µM) prior to the application of the pulse. When the membrane is intact, calcein is excluded from the interior of the GUVs, which appear dark in confocal cross sections. The fluorescence dye signal from the vesicle interior upon the application of a single strong DC pulse was used as a measure of prolonged membrane permeability. Calcein permeation in the vesicles was quantified by normalizing the internal fluorescence intensity inside a single GUV 5 minutes after macropore closure by the initial fluorescence intensity right after the macropore closure and the average external fluorescence intensity of the medium (SI section S7), thus eliminating contributions from differences in GUV size, background fluorescence and bleaching. No calcein was detected to flow inside the GUVs while macropores were open leaving the GUV interior black (Fig. 3A). GM1-free (pure POPC) GUVs remained impermeable to calcein 5 min after electroporation, indicating that pores in these membranes close completely and the membrane reseals. In contrast, asymmetric GM1-containing GUVs became permeable as observed by calcein leaking inside and the fraction of permeable GUVs increasing with GM1 fraction (Fig. 3). Prolonged permeation of asymmetric GM1-doped GUVs to calcein was also influenced by the degree of GM1 leaflet asymmetry as generated by different dilutions. Note that exploring entirely symmetric GM1-doped leaflets is not feasible because introducing calcein in the system requires the addition of the dye solution which is associated with small dilution (we avoided using extremely high calcein concentration, which could destabilize the vesicles). We explored 4 mol% GM1-doped GUVs, which were 1.25-, 3.33-and 10-fold diluted, and the number of leaky GUVs and overall leakage was observed to increase with dilution, i.e. with increasing GM1 leaflet asymmetry (Fig. S9). These results demonstrate that long-living submicroscopic pores are present in the GM1-doped asymmetric membranes.

**Fig. 3.**
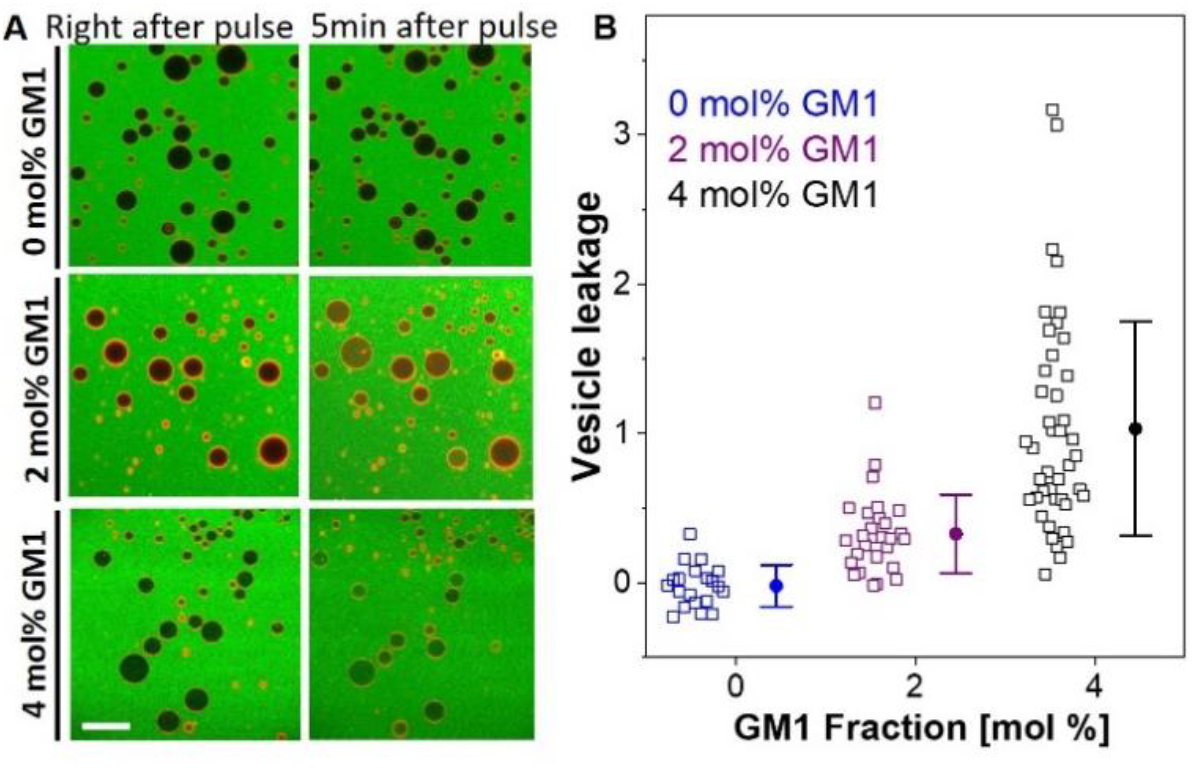
Long-term permeability of GUVs with increasing fractions of GM1. (A) Confocal images of GUVs illustrating calcein entry into GUVs with increasing fraction of asymmetric GM1 (0, 2 and 4 mol% top to bottom). The medium contained 5 µM calcein (green); the GUVs were labeled with 0.1 mol% DPPE-Rh (red). The snapshots in the first column were acquired ∼4 s after pulse application (0.6kV/cm, 3 ms) while the images in the second column show the same vesicles 5 minutes later. The scale bar is 100 µm. (B) Quantification of GUV leakage through fluorescence intensity analysis; see Eq. 3 in the SI for definition of leakage. Each open symbol corresponds to a measurement on a single GUV. Mean and standard deviation values are shown on the side.

Permeability has been observed to dramatically increase when approaching the main phase transition temperature of the membrane (52,53). Vesicles containing GM1 have been shown to exhibit gel-like domains (54) but at fractions higher than those examined here (above ∼5 mol%), which is why we can exclude this mechanism of increased permeability. The GM1-doped vesicles are also not leaky in the absence of electroporation (they preserve their sugar asymmetry and are impermeable to calcein over a period of at least 24 hours).

A plausible mechanism for stabilizing the submicroscopic pores causing long-term leakage could be steric obstruction by accumulated GM1 at the pore rims as well as protruding nanotubes. Indeed, GM1 bearing a single negative charge could be accumulated at the vesicle poles during the application of the pulse locally destabilizing the membrane as shown for vesicles with increasing surface charge (36).

## CONCLUSIONS

In summary, we used GM1-doped GUVs to mimic asymmetry of cell membranes. In particular, neuronal cells show elevated concentration of asymmetrically distributed GM1. We observed series of membrane remodeling effects resulting from the electroporation of GM1-containing asymmetric membranes, which all contribute to longer pore lifetimes and partial vesicle destabilization. When the GUV membrane is rendered asymmetric (by dilution), the desorption of GM1 from the outer leaflet of the vesicle membrane triggers the formation of inward tubes stabilized by negative spontaneous curvature. These tubes can physically obstruct the pores and membrane tubulation competes with pore closure, slowing down to the pore closure, reducing the effective membrane edge tension and rendering the membranes permeable at longer timescales. The decrease in edge tension depends on GM1 concentration and degree of leaflet asymmetry. Interestingly, pore lifetimes in the range of tens of seconds have been also observed in GM1-doped membranes but also in the presence of CTB (41), which forms homopentamers with GM1 (55) in the membrane, presumably leading to even slower reorganization of the pore. Our study also showed that increased fraction of GM1 stabilizes long-living submicroscopic pores and results in leaky vesicles after macropores close. Overall, our findings point to an additional role of GM1 in regulating the integrity of neuronal membranes (which are asymmetrically enriched in GM1) in lowering their stability under electrical perturbation and affecting membrane repair in wound healing.

## AUTHOR CONTRIBUTIONS

RD proposed the project. RD, RL and JS designed the experiments. MA, RL and JS performed the experiments and analyzed the data. All authors wrote the manuscript.

## DECLARATION OF INTEREST

The authors declare no competing interests.

## ACKNOWLEDGEMENTS

We thank R. Lipowsky, M. Miettinen and T. Bhatia for fruitful discussions on GM1-doped GUVs. M.A. thanks Y. Avalos-Padilla for help with analysis of fluorescence of CTB-Alexa binding to GM1. M.A. acknowledges funding from the International Max Planck Research School on Multiscale Bio-Systems.

## Notes

### Competing Interest Statement

The authors have declared no competing interest.

### Summary of Updates

- included statistics regarding the pore lifetimes of varying fractions of GM1 containing GUVs and compiled the data illustrating the scatter in a new panel in Fig. 1 - panel F - additional experiments to address effect of membrane asymmetry on vesicle leakage - data included in Fig. S9

