## Supporting information for "GM1 asymmetry in the membrane stabilizes pores"

#### S1. Preparation of giant unilamellar vesicles (GUVs)

GUVs were prepared by the electroformation method(1,2) as described in more detail in Refs.(3,4) Varying molar fractions of GM1 (Ovine Brain, Avanti Polar Lipids, Alabaster, AL) between 0 and 4 mol% in 1-palmitoyl-2-oleoyl-sn-glycero-3-phosphatidylcholine (POPC) (Avanti Polar Lipids) and 0.1 mol% 1,2-dipalmitoyl-sn-glycero-3-phosphoethanolamine-N-(lissamine rhodamine B sulfonyl) (DPPE-Rh) (Avanti Polar Lipids) were dissolved in dichloromethane:methanol (2:1) solution to a concentration of 4 mM. Both solvents were purchased from Sigma Aldrich, St. Louis, USA. Then, 20 µl of this lipid solution was spread as a lipid film on a pair of indium-tin oxide (ITO)-coated glass plates (PGO GmbH, Iserlohn, Germany), which are electrically conductive and preheated at 50 °C in the oven. A stream of $N_2$ was used to evaporate most of the solvent, and the plates were subsequently placed under vacuum at 50 °C for two hours to remove traces of the solvent. This temperature is above the main phase transition temperature of GM1 which is important for ensuring that the membrane is in the fluid and vesicle formation is favored.(5) For chamber assembly, a Teflon spacer of 2 mm thickness was placed between the ITO-glass plates and the chamber was filled with 1 mM HEPES buffer (pH 7.4, 0.5 Na HEPES; Sigma Aldrich) to hydrate the lipid film. Then, by applying a sinusoidal AC electric field at 10 Hz, electroswelling was initiated at 50 °C. The linear increment of the amplitude from 0.1 V (peak to peak) to 0.8 V (peak to peak) during the first 30 min, helping the swelling of the lipids, was followed by a constant application of 0.8 V for 90 min. Vesicle detachment from the glass surfaces was obtained by decreasing the voltage and frequency gradually to 0.5 V and 1 Hz over 60 min. GUVs were slowly cooled to room temperature over 30 min and then transferred to light-protective glass vials for storage at room temperature. The vesicles were used within 24 hours after preparation.

In order to assess the molar fractions of GM1 on symmetric GUVs, 25 µL of 10 mM Alexa Fluor 488 conjugated-cholera toxin B (CTB-Alexa) (Avanti Polar Lipids) were added to 5 mL of the vesicle solution to reach a final concentration of 50 nM and incubated for 20 min. For intact GUVs, one CTB binds to five GM1 molecules of the outer vesicle leaflet.(6) The fluorescence intensity of 0 – 4 mol% GM1 containing CTB-Alexa labelled GUVs were analyzed through confocal screenshots. The same microscope and objective settings were used during the quantification and generation of statistics for varying concentrations of GM1. Intensity measurements were performed through the "Radial profile extended plugin" from Philippe Carl installed from the ImageJ homepage. The radial intensity profile of GUVs was analyzed at the equatorial plane. A bigger centered circle was drawn on each analyzed vesicle to average the radial intensity in the circular area. An example of a confocal cross section and typical radial intensity profile of a 4 mol% GM1-doped GUV is plotted in Fig. S1A. The background fluorescence due to free CTB-

Alexa dye was subtracted during the data processing and then integrated peak area was taken as the normalized intensity for the comparison of GUVs with varying GM1 fractions. Fig. S1B demonstrates the resulting intensity averages of CTB-binding experiments for different GM1 fractions. The data shows a linear trend with the starting GM1 fractions in the membrane.

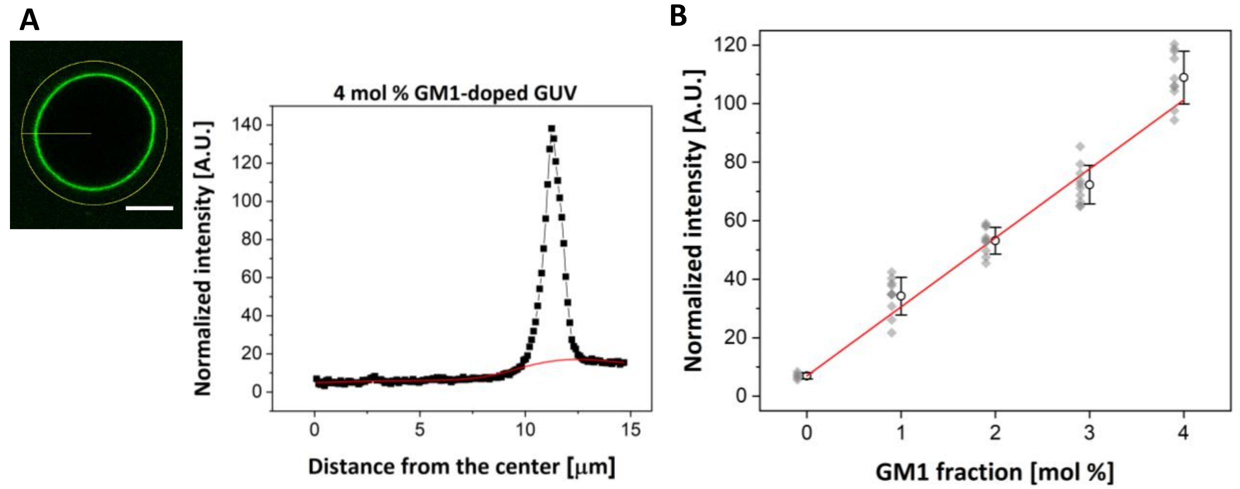


**Fig. S1.** The presence of GM1 on the outer leaflet of the vesicle membrane was confirmed from fluorescence of CTB-Alexa binding. (A) Representative confocal cross section and radial intensity profile of a symmetric GUV doped with 4 mol% GM1 analyzed via confocal microscopy and ImageJ radial profile analysis. The scale bar is 10 µm. Uniform fluorescence signal indicates homogeneous distribution of GM1 along the outer leaflet of the GUV. The graph shows the intensity peak which, after subtraction of the background (red curve) is integrated to quantify the fluorescence due to the binding of CTB-Alexa to the GM1 molecules in the outer leaflet of the GUV. (B) A plot of the resulting intensity averages of CTB-binding experiments for different GM1 fractions. For each GM1 fraction, 10 GUVs from 2 different preparations were analyzed. The data from individual GUVs are shown with light grey diamonds (overlapping single GUV experiments for same GM1 fraction appear as darker gray), while the mean values and standard deviations are displayed with black circles and line bars on the side. The red line shows the linear fit of the normalized mean intensity values for varying GM1 fractions.

**S2. Generation of asymmetric GUVs**

We generated GUVs with asymmetric distribution of GM1 in the two membrane leaflets. The asymmetry was established via GM1 desorption achieved with 10-fold dilution of the vesicle media with isotonic GM1-free 1 mM HEPES buffer, i.e. the buffer used for vesicle swelling. In order to visually assess the effects of asymmetry, pure POPC GUVs and 4 mol% GM1-doped GUVs were monitored with confocal laser scanning microscopy in non-diluted and 10-fold diluted conditions. In case of GM1-free vesicles (pure POPC), 100 GUVs in total from 8 independent samples were analyzed for each condition. The vesicles (with diameters above 10 µm) were inspected for defects with 3D confocal scans. In both diluted and non-diluted suspensions, 60 % of the GUVs appeared spherical and defect-free (Fig. S2A). The rest 40 % of the vesicles showed deformed shapes (dumbbell-like structure or tubulation) or had inner GUVs. For GM1-doped membranes, 120 GUVs containing 4 mol% GM1 from 10 different samples were examined. Before dilution, 80 % of the vesicles were defect-free. As a result of the 10-fold dilution with isotonic solution, 90 % of them formed inward tubes (Fig. S2B) stabilized from the negative spontaneous curvature of the asymmetric membrane. Some of the tubes appeared necklace-like with thickness in the suboptical range (Fig. S2C). Their morphology depends on the length and growth kinetics of the tubes, as well as on the elastic properties of GUVs.(3,7,8)

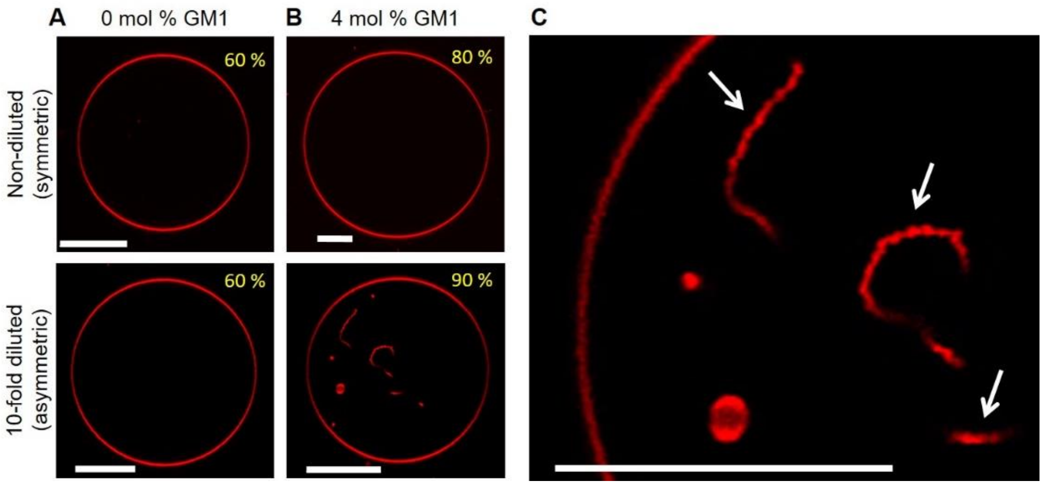


**Fig. S2.** GM1 desorption upon dilution of GUV external medium generates inward tubes stabilized by negative spontaneous curvature. (A) Confocal cross-sections of pure POPC in non-diluted and 10-fold diluted conditions. (B) Confocal cross-sections of 4 mol% GM1-doped GUV in non-diluted and 10-fold diluted conditions. In non-diluted conditions, both types of vesicles were smooth without inward or outward structures, i.e. with symmetric membranes. A 10-fold dilution in isotonic and GM1-free 1 mM HEPES solution caused the formation of internal nanotubes. The percentage of defect-free and tubulated vesicles relative to total GUV population is indicated in the upper-left corner of each snapshot. (C) Zoomed-in segment of the GM1-doped GUV in (B, lower panel) showing the necklace-like structure of the inward tubes (arrows). All scale bars are 20 μm.

### S3. GUV Electroporation and imaging

**Electroporation:** The GUVs were placed in a home-made electroporation chamber as described by Portet et al.(9) The chamber was assembled on a 26 x 56 mm cover glass equipped with a pair of parallel adhesive copper strips (3M, Cergy-Pontoise, France), 0.5 cm apart from each other. Then, small parafilm pieces were melted/glued onto the glass slide at a 1 cm distance from each other to seal the ends. An aliquot of 100 μL GUV solution was placed in the space between the copper tapes and a 22 x 22 mm coverslip (Thermo Fischer Scientific, Waltham, USA) was placed on top of the solution, forming a closed compartment. (Fig. S3A) Electrodes connected to βtech pulse generator GHT_Bi500 (βtech, l'Union, France) were attached to the edges of the copper tapes (Fig. S3B). The field strength and duration of the electric pulses were set to 150 – 200 V and 50 ms, respectively. In these pulse parameters, vesicles larger than 10 μm (as explored here) porate because the reached transmembrane potential exceeds the critical transmembrane potential for poration.(10,11) The response of the vesicles to a single (first) DC pulse was recorded for 2 – 5 minutes. The procedure was repeated on 10 – 15 GUVs for each composition.

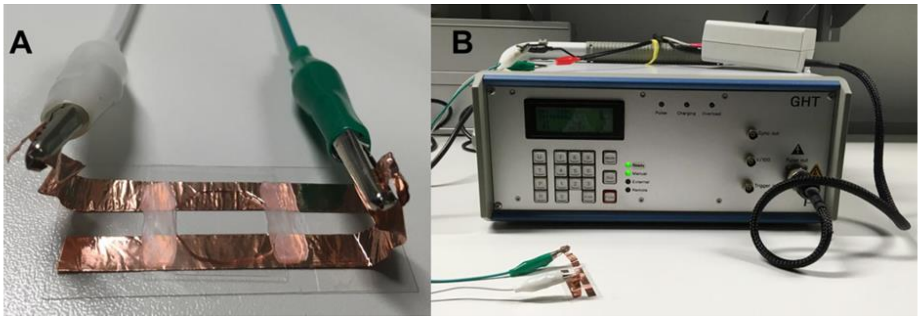


**Fig. S3.** Illustration of the electroporation set-up. (A) Home-made electroporation chamber. Two adhesive copper strips, 0.5 cm apart from each other were attached to a cover slide. Small parafilm pieces were placed orthogonally. After the addition of GUV solution, a small coverslip was placed on top and electrodes were connected to the copper strips. (B) Pulse Generator. A single DC pulse was applied (150 – 200 V and 50 ms) in each of the sample after which the sample was discarded.

**GUV imaging with confocal laser scanning microscopy:** The morphology of GUVs was monitored through Leica TCS SP5 or SP8 (Wetzlar, Germany) confocal microscopes. Observations were performed either via 40 x (0.75 NA) air or 63 x (1.2 NA) water immersion objectives. In the presence of the green dyes CTB-Alexa and calcein, excitation was performed with an argon laser line at 488 nm and the signal was collected in the range 495 – 550 nm. The red dye DPPE-Rh was excited with a diode-pumped solid-state laser at 561 nm and the emission was detected in the range 565 – 620 nm. The sequential mode of image acquisition was activated to minimize cross-talk. Images were recorded with 512 x 512 pixels and scanned with 400 Hz speed in the bidirectional mode with two line averages. In order to detect electroporation, a resonant scanner was used, which increased the scanning speed to 800 Hz. The images could be obtained on both forward and back scans, allowing to record 36 frames per second (fps). Image analyses were performed through Leica LASX software (Jena, Germany) and ImageJ (NIH, USA). Analyzed data were plotted through Origin Pro software.

**Epifluorescence microscopy and high-speed imaging:** Edge tension analysis of GUVs during the electroporation experiments was conducted with Axio Observer D1 epifluorescence microscope (Zeiss, Germany) equipped with a sCMOS camera (pco.edge, PCO AG, Kelheim, Germany). Images were recorded in epifluorescence mode with a 40 x / 0.60 Korr Ph2 objective (Zeiss) at a varying acquisition speed between 50-100 fps. The samples were irradiated by using the HBO 100 W mercury lamp of the microscope in epi-illumination mode. For red irradiation, the light from the mercury lamp passed through 560 / 40, 585, and 630 / 75 nm filter cube, respectively. The image analyses were performed with the ImageJ software.

**Phase contrast microscopy and high-speed imaging:** For phase contrast microscopy, GUVs were imaged by a Zeiss Axiovert 200 microscope (Jena, Germany) equipped with an ultra-high-speed digital camera v2512 (Phantom, Vision Research, New Jersey, USA). A halogen lamp HAL 100 was used for the illumination of the samples and all the images were collected with a 40x Ph2 objective at acquisition speed of 5000 fps.

### S4. Membrane tubulation after pore formation

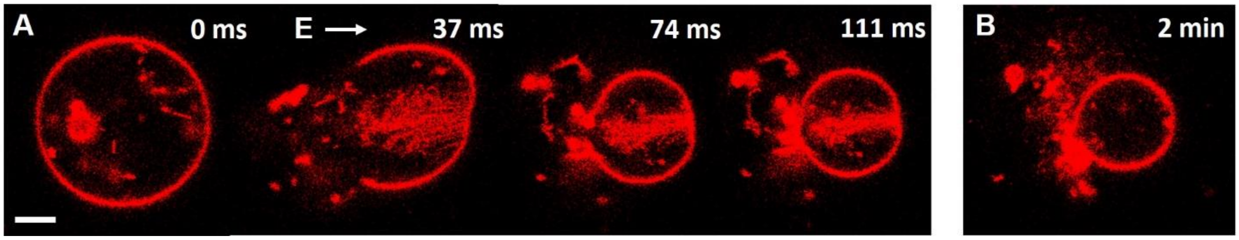


**Fig. S4.** Sprouting of membrane nanotubes from 4 mol% GM1-doped asymmetric GUV; also shown in Movie S3. (A) A sequence of images obtained with confocal microscopy during and after application of a DC pulse (0.3kV/cm, 50 ms). The direction of the electric field is illustrated with the arrow. The time is relative to the beginning of the pulse and shown on the left-upper side of each snapshot. The membrane contains 0.1 mol% DPPE-Rh and the GUV was 10-fold diluted with 1 mM isotonic HEPES buffer. (B) Cross section of the vesicle 2 minutes after the pulse application. The scale bar is 10 μm.

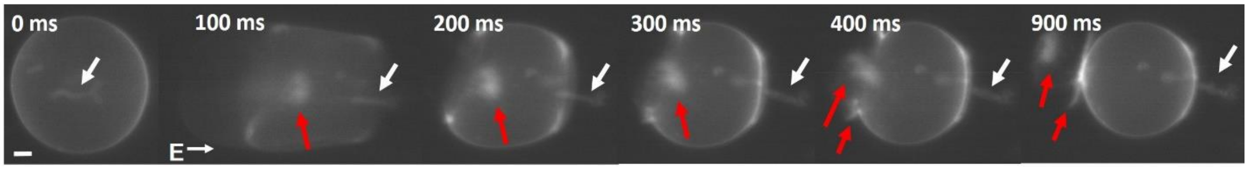


**Fig. S5.** Sprouting of membrane nanotubes from 4 mol% GM1-doped asymmetric GUV; also shown in Movie S4. A sequence of images obtained with epifluorescence microscopy during and after the application of a DC pulse (0.3kV/cm, 50 ms). The direction of the electric field is illustrated with the arrow. The time is relative to the beginning of the pulse and shown on the left-upper side of each snapshot. The white arrow points to an inward tube present already before the application of the pulse. Red arrows point to tubes protruding inside the vesicle and at the pore rims formed after the application of the pulse and expelled outside the vesicle after the application of the pulse. The membrane contains 0.1 mol% DPPE-Rh and the GUV was 10-fold diluted with 1 mM isotonic HEPES buffer. The scale bar is 10 μm.

### S5. Pore stability

We compare the edge energy to the bending energy of the asymmetric membrane as the pore radius increases. When the membrane bending energy is larger than the open pore edge energy, it is unfavorable to close the pore and the vesicle is destabilized whereby the flat membrane would reshape into tubular structure rather than closing the pore (see Fig. S6).

The energy of a porated vesicle of radius $R$ is a sum of the Helfrich and the rim energy:

$$E = 2\kappa \int_A (M-m)^2 dA + 2\pi\gamma r, \qquad \text{(S1)}$$

where $\kappa$ is the membrane bending rigidity, $M = 1/R$ is the vesicle mean curvature, $m$ is the membrane spontaneous curvature (assumed constant), $\kappa_G$ is the Gaussian curvature modulus, $r$ is the pore radius and $\gamma$ is the edge tension. The integral in Eq. S1 is over the vesicle (non-porated) area $A = 2\pi R^2\left[1+\sqrt{1-(r/R)^2}\right]$. Here, we have ignored Gaussian curvature contributions because the topology of a vesicle with expanding pore is preserved.

For a pore to have the tendency to expand, as is the case of asymmetric vesicles, the energy of the system should decrease, i.e.

$$dE/dr = -\frac{4\pi\kappa R(1/R-m)^2}{\sqrt{(R/r)^2-1}} + 2\pi\gamma < 0\ . \qquad \text{(S2)}$$

Above we assumed that the vesicle radius remains constant, which for small changes in the pore radius is roughly the case. Equation S2 implies that the edge tension term should be dominated by curvature contributions, implying that the tension associated with the spontaneous curvature (spontaneous tension) can act to pull the pore open.

The inequality S2 can be transformed into

$$r > \left(\frac{4\kappa^2 m^4}{\gamma^2} - \frac{1}{R^2}\right)^{-1/2} \qquad \text{(S3)}$$

Considering the large size of the explored GUVs ($R$ is typically above 20 µm), the second term in the brackets can be ignored, leading to a critical pore size, $r_c$, above which the pore would rather expand and the vesicle will collapse

$$r > r_c \equiv \gamma/2\kappa m^2 \qquad \text{(S4)}$$

Introducing the values $\kappa = 1.2\times10^{-19}$ J and $\gamma$ = 17 pN (see Fig. 2C in the main text) and using $m = -1/460$ $nm^{-1}$ for vesicles prepared with 2 mol% GM1 and $\gamma$ = 7 pN and $m = -1/220$ $nm^{-1}$ for vesicles prepared with 4 mol% GM1 (3) we obtain for the critical pore size $r_c$ ~15 µm and 1.4 µm for the respective compositions.

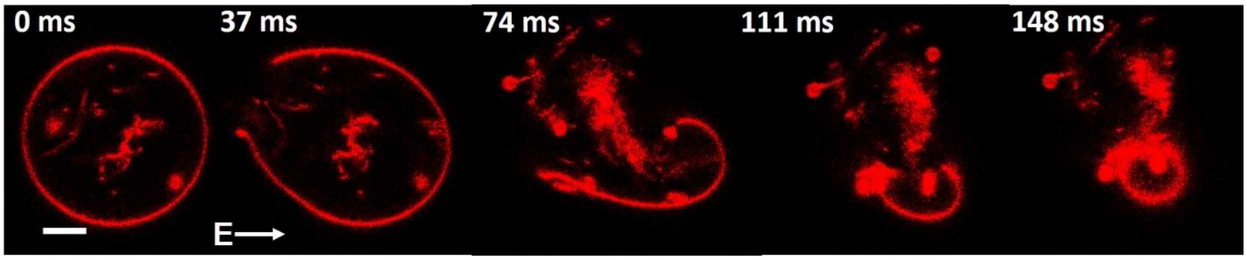


**Fig. S6.** Complete transformation of 4 mol% GM1-doped asymmetric GUV to a tubular network. A sequence of images obtained with confocal microscope during and after application of a DC pulse (0.3 kV/cm, 50 ms). The direction of the electric field is illustrated with the arrow. The time is relative to the beginning of the pulse and shown on the left-upper side of each snapshot. The membrane contains 0.1 mol% DPPE-Rh and the GUV was 10-fold diluted with 1 mM isotonic HEPES buffer. The scale bar is 10 μm.

### S6. Edge tension analysis of GUVs

Edge tension measurements are based on the theory described by Brochard-Wyart et al.(12,13) In order to study in more details the effect of GM1, we measured edge tension for GUVs with increasing GM1 molar fraction. To facilitate the visualization of membrane pores and still have a good temporal resolution, pore closure dynamics were recorded using epifluorescence microscopy. The data were evaluated by manually measuring the pore radius, $r$, from every recorded image. Pore dynamics are characterized by the quantity, $R^2 ln(r)$ as a function of time, in which $R$ and $r$ are GUV and pore radius, respectively. Three typical datasets for the different GM1 fractions are shown in Fig. 2C. The linear behavior of the data and the precision of the values of the slopes in the graph for the same GM1 fractions validated the accuracy and the consistency of the experiments with the theoretical model. The straight solid lines represent the linear fits of the quasi-static leakout regime. The edge tension is obtained from the slope of linear fit, $\alpha$, using the equation of $\gamma = -\left(\frac{3}{2}\right)\pi\eta\alpha$, where $\gamma$ and $\eta$ are the edge tension and the medium viscosity ($\eta$ = 0.89×10$^{-4}$ Pa.s for our experimental conditions), respectively. The data differences in the y-axis result from the differences in the sizes of the GUVs. GUVs with the same GM1 compositions have different absolute values but similar slopes, which is indicative for similar edge tensions. The obtained values for 0, 2 and 4 mol% of GM1-doped GUVs in non-diluted and 10-fold diluted conditions are displayed in Fig. S7 and the mean values of the edge tensions for each composition are listed in Table S1. Standard error values for the mean edge tension measurements of each fraction are also found as reasonable (less than 15%), indicating the precision of our experiments.

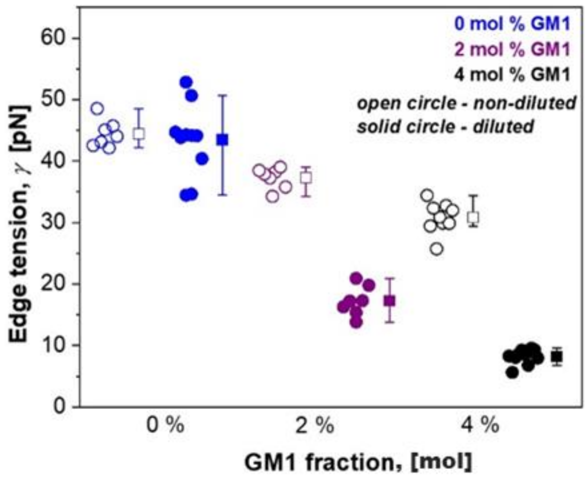


**Fig. S7.** Edge tension values for vesicles with different molar fractions of GM1 in non-diluted (open circles) and 10-fold diluted (solid circles) systems. Each circle indicates a measurement of an individual GUV. Mean and standard deviation values are shown by the squares and the line bars, respectively. Blue, purple and black corresponds to 0, 2, 4 mol% of GM1 fractions (same data as in Fig. 2C in the main text). The results indicate that increasing GM1 fraction and asymmetry decrease membrane edge tension.

**Table S1.** Mean edge tension values for the GUVs with varying GM1 fraction

| **GM1 fraction (mol% )** | **Edge tension (pN)** | |
|---|---|---|
| | **Non-diluted (symmetric)** | **Diluted (asymmetric)** |
| 0 | 44.4 ± 2.2 | 43.3 ± 5.6 |
| 2 | 37.2 ± 1.7 | 17.2 ± 2.3 |
| 4 | 30.8 ± 2.5 | 8.2 ± 1.1 |

### S7. Analysis of GUV prolonged permeability

The prolonged permeability of GM1-doped asymmetric GUVs after the macropore closure was firstly observed under phase contrast imaging. GM1-doped asymmetric GUVs tended to lose their optical contrast 15 minutes after the macropore closure (Fig. S8).

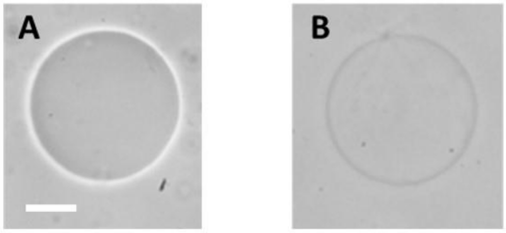


**Fig. S8.** Long-term permeation of 4 mol% asymmetric GM1-doped GUV right after the macropore closure (A) and 15 minutes after macropore closure (B). The GUV prepared in 200 mM sucrose was 10-fold diluted in isotonic glucose solution. The contrast loss of the GUV was monitored via phase contrast microscopy after the application of a DC pulse with amplitude of 0.6 kV/cm and duration of 2 ms. The scale bar is 10 µm. Due to the different refractive index of the sugar solutions, the vesicles look dark before poration and after the macropore closure. However, during prolonged observations, 15 minutes after the macropore closure, GUV had lost its optical contrast implying that submicroscopic pores remain present.

The long-term permeability of GUVs upon a DC pulse application was quantified by using the small water-soluble marker calcein. An aliquot of 30 µL of GUVs prepared in 200 mM sucrose solution (200 mosmol/kg) was diluted with 65 µL isotonic glucose solution and then 5 µL of 0.1 mM calcein was added to the diluted system. The sample was incubated for 15 minutes to ensure homogeneity. Then, a single DC pulse with 0.5 – 0.6 kV/cm amplitude and 3 ms duration was applied. Long-term GUV permeation was assessed 5 minutes after the application of the pulse from confocal cross section images of the vesicles in bulk. The calcein influx in GUVs containing 0 − 4 mol% of GM1 was measured from the change in the fluorescence intensity over time in the GUV interior assessing a circular ROI of nominal radius $r_n$ = 5 µm. The fluorescence inside a GUV 5 minutes after the macropore closure was normalized with respect to the fluorescence just after macropore closure and correcting for the fluorescence of the external GUV medium. The GUV leakage, $L$, was defined as:

$$L = \frac{\left(\frac{F_{in}}{F_{out}}\right)_5 - \left(\frac{F_{in}}{F_{out}}\right)_0}{\left(\frac{F_{in}}{F_{out}}\right)_0} \tag{3}$$

where $F_{in}$ is the average fluorescence intensity inside the GUV and $F_{out}$ is the average fluorescence of the medium outside the GUV; the subscripts "5" and "0" refer to the ratios measured 5 minutes after the macropore closure and right after the macropore closure, respectively. This definition accounts for fluorescence fluctuations due to measurements at different height in the sample (i.e. different vesicle size) or local changes of fluorescence.

The resulting mean intensity values with increasing GM1 fraction were plotted using Origin Pro software. The statistics were generated over 20 − 40 GUVs obtained from 3 independent samples. The analyzed GUVs ranged in diameter between 25 and 100 µm.

In order to probe the effects of degree of asymmetry on the long-term permeation of GM1-doped GUVs after the exposure to a DC pulse, aliquots of 4 mol% GM1 GUVs grown in 200 mM sucrose were diluted at different dilution ratios with isotonic 200 mM glucose solution and the leakage of 20 – 40 vesicles were analyzed for calcein permeation 5 min after macropore closure as explained above. Prior to electric field application, the GUVs did not show any permeation to calcein for 24 hours. After the application of electric field and the closure of macropores, more diluted samples (i.e. with more asymmetric membranes) showed higher number of and more leaky GUVs, indicating that increasing the degree of GM1 asymmetry in the membrane leaflets results in more pronounced calcein permeation and higher proneness to the formation of long-living sub-microscopic pores (Fig. S9).

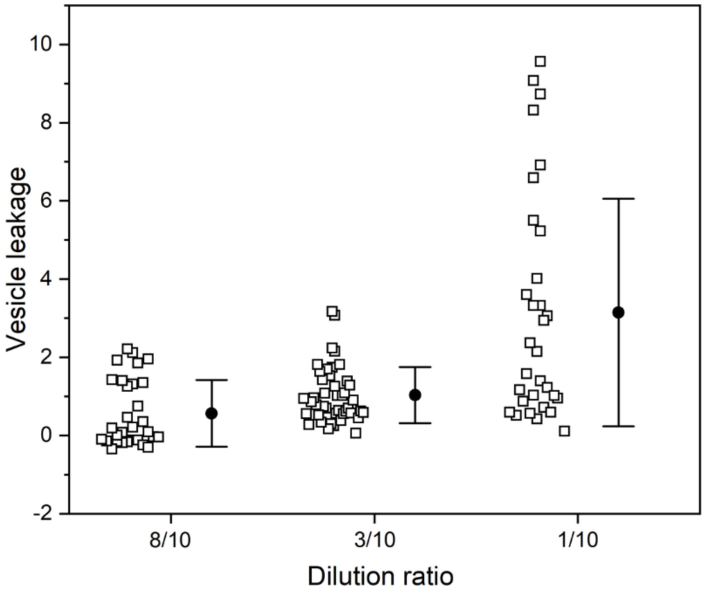


**Fig. S9.** Long-term calcein permeation of 4 mol% GM1 containing GUVs with different dilution ratio i.e. membrane asymmetry. The external medium contained 5 µM calcein; the GUVs were labeled with 0.1 mol% DPPE-Rh, grown in 200 mM sucrose solution and diluted in 200 mM glucose solution. Dilution ratios in terms of volumes of GUV suspension to glucose solutions were 8:2, 3:7 and 1:9 corresponding respectively to 1.25-fold (10/8), 3.33-fold (10/3) and 10-fold dilutions. GUVs were exposed to 0.6 kV/cm, 3 ms DC pulse and calcein permeation was monitored 5 min after the pulse application. GUV leakage was quantified via fluorescence intensity analysis; see Eq. 3 for definition of leakage. Each open square corresponds to a measurement on a single GUV. Mean and standard deviation values are shown on the side. At 0.05 level, the mean values of each dilution ratio are significantly different from each other based on one-way ANOVA testing and paired-sample t-test ($p<0.05$).

**Supplementary movies**

**Movie S1.** A slowed down video of a typical electroporation process of a 4 mol% GM1-doped asymmetric GUV corresponding to the image sequence shown in Fig. 1E in the main text (where images were rotated to keep the same direction of the field in all figures). The sequence was imaged in epifluorescence during and after application of a DC pulse (0.3 kV/cm, 50 ms). The time is relative to the beginning of the pulse and shown on the left-upper side of each snapshot. The membrane contains 0.1 mol% DPPE-Rh and the GUV was 10-fold diluted with 1 mM isotonic HEPES buffer. Video was processed with Fiji and slowed down to 8 frames per second.

**Movie S2.** A slowed down video of a typical electroporation process of a 4 mol% GM1-doped asymmetric GUV (same vesicle as in Fig. 2 in the main text), presenting a sequence of confocal microscopy images acquired during and after application of a DC pulse (0.3kV/cm, 50 ms). The time is relative to the beginning of the pulse and shown on the upper left corner of each snapshot. The membrane contains 0.1 mol% DPPE-Rh and the GUV was 10-fold diluted with 1 mM isotonic HEPES buffer. The video was processed with Fiji and slowed down to 4 frames per second.

**Movie S3.** A slowed down video showing membrane nanotubes sprouting and being expelled from a 4 mol% GM1-doped asymmetric GUV (corresponding to the image sequence shown in Fig. S4 where images were rotated), representing a time sequence of confocal microscopy images during and after application of a DC pulse (0.3kV/cm, 50 ms). The time is relative to the beginning of the pulse and shown on the left-upper side of each snapshot. The membrane contains 0.1 mol% DPPE-Rh and the GUV was 10-fold diluted with 1 mM isotonic HEPES buffer. Video was processed with Fiji and slowed down to 3 frames per second.

**Movie S4.** A slowed down video of sprouting of membrane nanotubes from 4 mol% GM1-doped asymmetric GUV (corresponding to the image sequence shown in Fig. S5 where images were rotated). A sequence of images obtained in epifluorescence during and after application of a DC pulse (0.3 kV/cm, 50 ms). The time is relative to the beginning of the pulse and shown on the left-upper side of each snapshot. The membrane contains 0.1 mol% DPPE-Rh and the GUV was 10-fold diluted with 1 mM isotonic HEPES buffer. Video was processed with Fiji and slowed down to 3 frames per second.

**Movie S5.** A slowed down video of bursting of 4 mol% GM1-doped asymmetric GUV to a tubular network shown in Fig. S6. A sequence of images obtained with confocal microscope during and after application of a DC pulse (0.3kV/cm, 50 ms). The time is relative to the beginning of the pulse and shown on the left-upper side of each snapshot. The membrane contains 0.1 mol% DPPE-Rh and the GUV was 10-fold diluted with 1 mM isotonic HEPES buffer. Video was processed with Fiji and slowed down to 3 frames per second.
